# Genome-scale mapping reveals complex regulatory activities of RpoN in *Yersinia pseudotuberculosis*

**DOI:** 10.1101/2020.08.19.258541

**Authors:** F.A.K.M. Mahmud, K. Nilsson, A. Fahlgren, R. Navais, R. Choudhry, K. Avican, M. Fallman

## Abstract

RpoN, an alternative sigma factor commonly known as sigma 54, is implicated in persistent stages of *Yersinia pseudotuberculosis* infections in which genes associated with this regulator are upregulated. We here combined phenotypic and genomic assays to provide insight into its role and function in this pathogen. RpoN was found essential for *Y. pseudotuberculosis* virulence in mice, and *in vitro* functional assays showed that it controls biofilm formation and motility. Mapping genome-wide associations of *Y. pseudotuberculosis* RpoN using chromatin immunoprecipitation coupled with next-generation sequencing identified an RpoN-binding motif located at 103 inter- and intragenic sites on both sense and anti-sense strands. Deletion of *rpoN* had a large impact on gene expression, including down-regulation of genes encoding proteins involved in flagellar assembly, chemotaxis, and quorum sensing. There were also clear indications of cross talk with other sigma factors, together with indirect effects due to altered expression of other regulators. Matching differential gene expression with locations of the binding sites implicated around 130 genes or operons potentially activated or repressed by RpoN. Mutagenesis of selected intergenic binding sites confirmed both positive and negative regulatory effects of RpoN binding. Corresponding mutations of intragenic sense sites had less impact on associated gene expression. Surprisingly, mutating intragenic sites on the anti-sense strand commonly reduced expression of genes encoded by the corresponding sense strand.

**IMPORTANCE:** The alternative sigma factor, RpoN (σ 54), which is widely distributed in eubacteria have been implicated to control gene expression of importance for numerous functions including virulence. Proper responses to host environments are crucial for bacteria to establish infection and regulatory mechanisms involved are therefore of high interest for development of future therapeutics. Little is known about the function of RpoN in the intestinal pathogen *Y. pseudotuberculosis* and we therefore investigated its regulatory role in this pathogen. This regulator was indeed found to be critical for establishment of infection in mice, likely involving its requirement for motility and biofilm formation. The RpoN regulon involved both activating and suppressive effects on gene expression which could be confirmed with mutagenesis of identified binding sites. This is the first of its kind study of RpoN in *Y. pseudotuberculosis* revealing complex regulation of gene expression involving both productive and silent effects of its binding to DNA providing important information about RpoN regulation in enterobacteria.

## INTRODUCTION

During infection of a host, bacteria are exposed to rapid changes in the environment, such as changes in temperature, pH, osmolarity, nutrient levels, and immune cell attacks. Bacteria usually cope with these types of changes through stress responses, alterations of their gene expression that are adaptive to the new environment (1). This type of infection-associated transcriptomic reprogramming was obvious in a previous *in vivo* transcriptomic study of *Y. pseudotuberculosis* isolated from cecal lymphoid compartments of infected mice (2). In that model, plasmid-encoded virulence genes known to be necessary for tissue invasion and resistance towards initial attacks from phagocytes were highly expressed during the early phase of the infection. After about one-and-a-half months of symptomless infection, the expression pattern had changed so that genes encoding proteins involved in adaption and resistance to different types of stresses dominated, while expression of the plasmid-encoded virulence genes was considerably reduced. This highlights the importance for bacteria of both adapting to new environments and the regulatory mechanisms involved. Hence, increasing our understanding of bacterial function during infection is of great interest. Mechanisms of bacterial adaptation inform choices of potential targets for new antibiotics, with gene products required to maintain infections considered more promising than those of classical virulence genes.

Transcriptional reprogramming is commonly controlled by various transcriptional regulators that are activated in response to external signals. A major class of transcriptional regulators are sigma factors, which upon activation associate with the core RNA polymerase (RNAP), promoting its binding to specific initiation sites and subsequent open complex formation for transcription of downstream genes (3). There are different types of sigma factors in bacteria, where RpoD or sigma70 (σ^70^) is the primary and house-keeping sigma factor active during exponential growth (4-6). Other, alternative sigma factors such as RpoE, RpoS, RpoH, and RpoN, which recognize promoter sequences distinct from that of RpoD, regulate transcription during specific conditions, allowing expression of genes required for the bacteria to cope with and adapt to particular situations (5). One of the sigma factors that attracted interest during the analysis of data from our previous *in vivo* transcriptomic analysis of *Y. pseudotuberculosis* was RpoN or sigma 54 (σ^54^), which together with many of its associated proteins, including activators and modulating proteins, was upregulated during the persistent stages of infection, when the expression of genes important for adaptation to the tissue environment dominated (2). RpoN has been reported to control regulation of genes involved in nitrogen metabolism, flagella, and motility, but biofilm formation and quorum sensing can also be affected in *rpoN* mutant strains (7-10). In some species, RpoN also influences regulation of Type III and Type VI secretion (8, 11-13), and there are many reports implicating RpoN as a regulator of bacterial virulence (9, 14, 15).

RpoN is structurally and functionally distinct from other sigma factors in that transcription initiation commonly depends on its binding to activating proteins termed bacterial enhancer-binding proteins (EBPs) (6). These EBPs use ATP catalysis to remodel RNAP DNA binding to initiate transcription (16). The affinity of RpoN for the core RNAP is higher than for most other RpoD-related alternative sigma factors, allowing it to compete efficiently for RNAP binding. Regulation by RpoN can be either direct or indirect via activation of different positive or negative regulators, including other sigma factors (5, 17, 18). Compared with other sigma factors, direct cross talk whereby different sigma factors regulate the same gene is particularly high for RpoN and is also commonly seen for genes that encode proteins involved in complex processes with different levels of regulation, such as adaption, chemotaxis, adhesion, and protein secretion (19). Adding to the complexity is the variation between different bacteria in the specific signals regulating RpoN and the specific downstream outcomes. One example is biofilm formation, where a deletion of the *rpoN* gene results in severe effects on the capacity to form biofilms in many bacteria, but where the opposite is seen in some other bacteria (20, 21).

This study investigated the regulatory role of RpoN in *Y. pseudotuberculosis*, which had not previously been addressed in detail. Neither has the RpoN regulon been defined for this pathogen. The results revealed that RpoN is crucial in *Y. pseudotuberculosis* to establish infection and is required for biofilm formation and motility. Chromatin immunoprecipitation coupled with next-generation sequencing (ChIP-seq) was used to determine genome-wide binding and revealed more than 100 RpoN binding sites with both inter- and intragenic locations. Transcriptomic data from bacteria lacking *rpoN* implied a complex regulatory network with direct or indirect effects. Matching the locations of ChIP peaks with transcriptomic data allowed retrieval of more than 130 genes potentially regulated by RpoN, some novel and some known from previous studies. Mutagenesis of selected RpoN binding sites confirmed both activating and suppressive roles of upstream intergenic RpoN-binding. This was not seen for sites of intragenic binding to the sense strand. In contrast, mutation of RpoN binding motifs on the anti-sense strand commonly resulted in suppressed expression of the gene on the sense strand, implicating a novel regulatory mechanism.

## RESULTS AND DISCUSSION

### RpoN participate in regulation of *Y. pseudotuberculosis* biofilm formation and motility and is required for virulence

To reveal the importance of RpoN in *Y. pseudotuberculosis* virulence, mouse infection studies were conducted using the *Y. pseudotuberculosis* WT strain as well as an *rpoN* deletion mutant strain. In accordance with prior results in this virulence model for *Y. pseudotuberculosis* acute infection, the WT strain initially colonized all infected mice, and all showed clear signs of disease and succumbed at day 5–7 post infection (Fig. 1A and Fig. S1A). The Δ*rpoN* strain on the other hand, had colonized all mice to a low extent one day after infection and showed increased colonization at day 3, but then the infection declined and the majority were cleared at day 14–21. At 29 days post infection, all mice had cleared the infection (Fig. 1A and Fig. S1A). None of the mice infected with the *ΔrpoN* strain showed signs of disease during the infection period, indicating clear attenuation. We also employed the *Y. pseudotuberculosis* infection model for persistent infection of the cecal lymphoid compartment, where a fraction of mice infected with low doses of the WT strain carry bacteria in cecal tissue for prolonged times without showing symptoms of disease (22). Upon low-dose infection with the Δ*rpoN* strain, only 75% of infected mice were initially colonized compared with mice infected with the corresponding WT strain, all of which were colonized (Fig S1B). This suggest a critical role for this regulator during the early stages of infection. We next tested the Δ*rpoN* strain in different phenotypic assays and found that the mutant strain was deficient in biofilm formation and motility (Fig. 1C and D). The biofilm and motility phenotypes could be complemented by expressing RpoN in *trans* (Fig. 1C and D). Hence, it is obvious that the *ΔrpoN* mutant strain has limited functional capacity that likely contributes to the observed attenuation in virulence. Together, these data indicate a pivotal role for RpoN in *Y. pseudotuberculosis* virulence.

**FIG 1.**
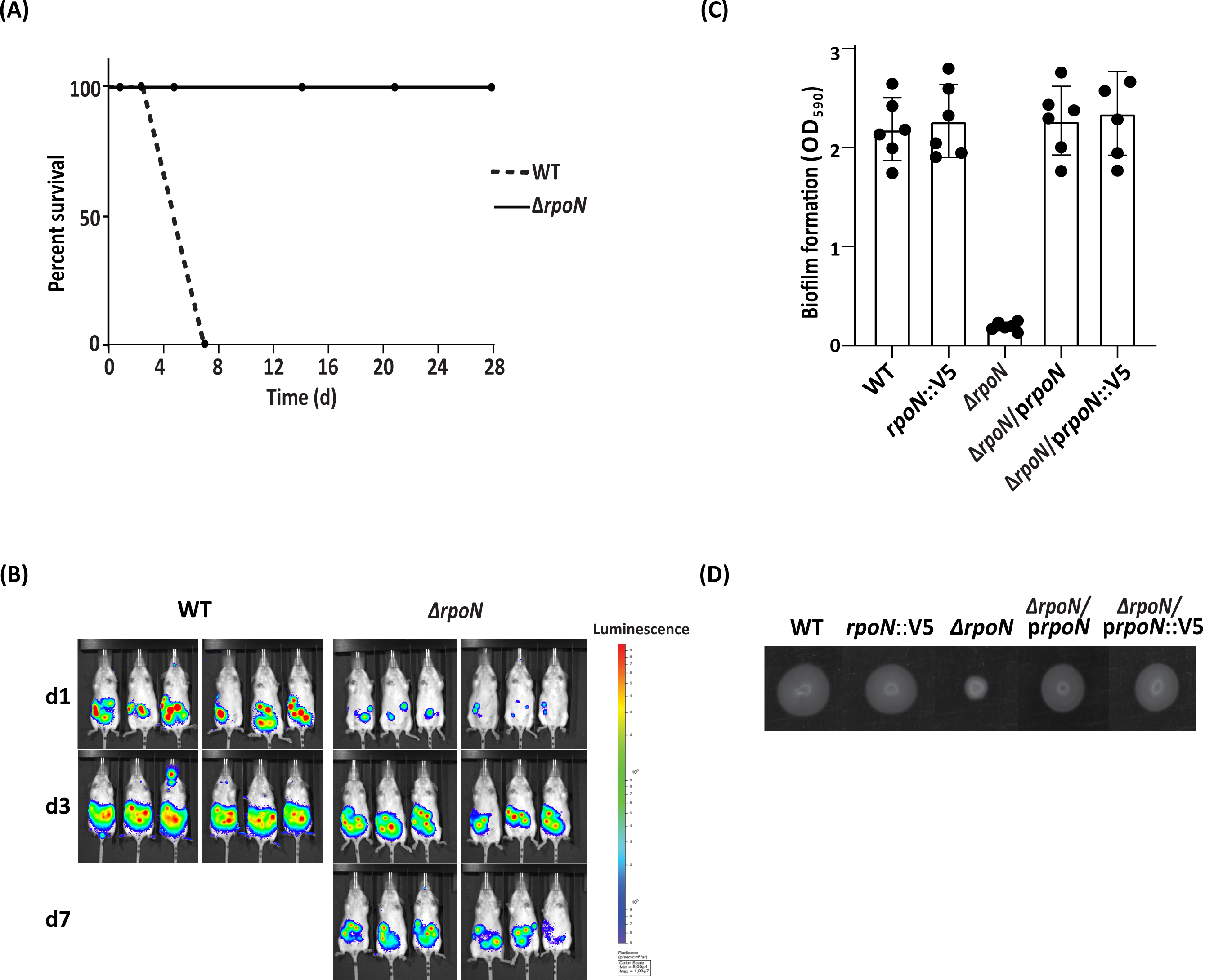
RpoN is required for *Y. pseudotuberculosis* virulence and essential for motility and biofilm formation. (A) Female FVB/N mouse (n = 6) were infected with 4.2 × 108 CFU/ml of *Y. pseudotuberculosis* WT (dashed line) or the isogenic Δ*rpoN* mutant strain (solid line). Data are presented as percent live mice during the course of infection (0–28 days). (B) *In vivo* images showing anesthetized mice during the first 7 days of infection. Colors indicate levels of light emitted by bioluminescent bacteria. Mice infected with *Y. pseudotuberculosis* WT become sick at day 5–7 and were sacrificed. (C) Biofilm formation by *Y. pseudotuberculosis* WT, and the corresponding strains with *rpoN*::V5 inserted in the chromosome, Δ*rpoN*, Δ*rpoN* complemented *in trans* with p*rpoN*, and Δ*rpoN* complemented *in trans* with p*rpoN*::V5. Biofilm mass was determined by dissolving biofilm material and measuring the OD_590_. Data are presented as means ± SEM for six individual experiments (D) Motility in agarose by the same strains as indicated in B. One representative experiment of three replicates is shown.

### RpoN binds to multiple sites on the *Y. pseudotuberculosis* chromosome

Since previous studies of RpoN-mediated events have indicated variation in the regulatory network between different bacterial species, and since studies of RpoN in *Y. pseudotuberculosis* are limited, we set out to uncover the regulatory mechanisms involved. As a first step we employed ChIP-seq to determine the RpoN DNA-binding sites in *Y. pseudotuberculosis*. ChIP-seq strongly depends on an appropriate antibody, and we chose to fuse the C-terminus of RpoN to the 3×V5 epitope, which has been shown to function in similar approaches in other bacteria (23). We generated a vector construct to overexpress RpoN-V5 (p*rpoN*::V5) and also a strain expressing RpoN-V5 from its native promoter, by inserting the sequence for the V5-tag in the 3’end of the *rpoN* gene (*rpoN*::V5). Repeating the assays for biofilm formation and motility and including the Δ*rpoN/*p*rpoN*::V5 and the *rpoN*::V5 strains showed that the V5 tag did not interfere with RpoN function (Fig. 1C and D). It is commonly assumed that RpoN levels and binding to RNAP and to DNA are relatively stable and that the EBPs play the regulatory role. However, our finding of differential expression of *rpoN in vivo* compared with its expression level *in vitro* (2), and the possibility of chromosomal structural changes influencing gene expression under certain conditions (24, 25), prompted us to map the binding of RpoN in bacteria subjected to more than one condition. The conditions used were exponential growth at 26°C, where RpoN-V5 either is over-expressed (p*rpoN*::V5) in *trans* or in *cis* (*rpoN*::V5), the latter to ensure proper stoichiometry and thereby avoid side effects of competition with other sigma factors. We also included samples subjected to virulence-inducing conditions, which includes a shift to 37°C and depletion of extracellular Ca^2+^ for 75 minutes that results in expression of the virulence plasmid (26) in the *rpoN*::V5 strain.

The resulting ChIP-seq data were subjected to a high-stringency bioinformatic analysis with a cut-off of a 2.5-fold difference over genomic noise. This identified 119 ChIP-seq peaks representing putative sites for RpoN binding. The number of peaks was in the range previously shown for *E. coli, Salmonella* Typhimurium, and *V. cholerae* (8, 27, 28). Some of the peaks were narrow and distinct, covering 200–300 nucleotides, whereas others were relatively broad and covered 300–800 nucleotides (Table S1). The predicted peaks were used to identify a common sequence motif in the ± 50 bp regions of the peak center. We identified a motif that resembled the RpoN −24/12 promoter element found in other bacteria (27, 29, 30) (Fig. 2A). To determine motif strength, NN-GG-N9-TGC-NN was used as base for position-weight matrix calculations. The identified motifs had PSSM (position-specific scoring matrix) scores ranging from 3 to 12, and the motif sequences were positioned 20 bp upstream of the peak center (Fig. 2B).

**FIG 2.**
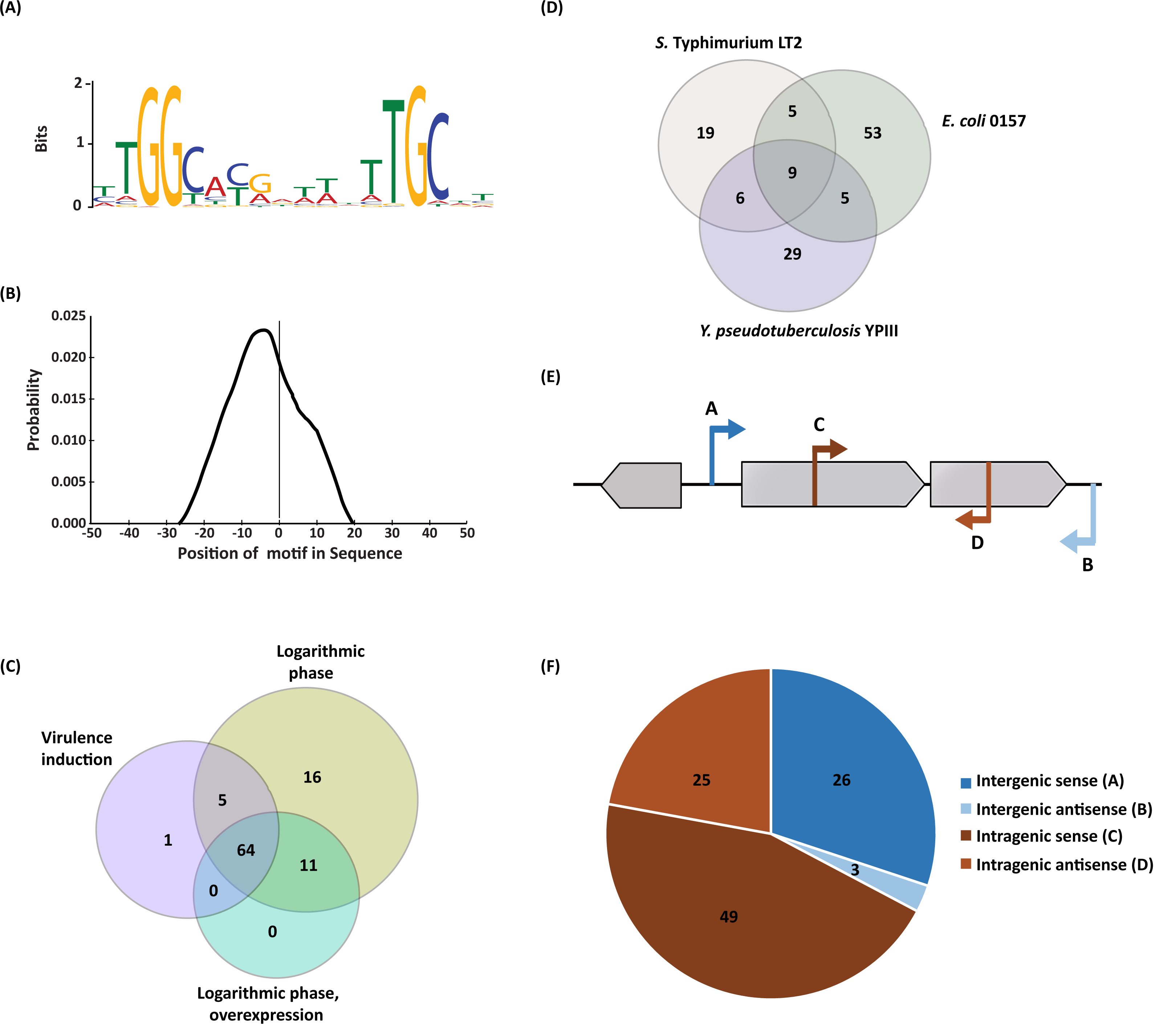
ChIP-seq identifies RpoN binding sites in *Y. pseudotuberculosis* genome. (A) RpoN consensus binding motif in *Y. pseudotuberculosis* derived from 119 ChIP-seq peaks, determined using BCRANK (E-value = 1.8e–110). (B) Position of motifs relative to peak centers calculated using the Centrimo tool, MEME suite. The graph indicates the average density of motif position for all 119 motif-containing regions, using 10 bp bins from position – 50 to +50 relative to the ChIP-seq peak. (C) Venn diagram showing the number of RpoN binding sites identified by ChIP-sequencing of *Y. pseudotuberculosis* in different growth conditions. (D) Venn diagram showing number of RpoN binding sites with homologous genomic positions in *Y. pseudotuberculosis, E. coli*, and *S*. Typhimurium. Those genes which were found to be conserved based on function and phylogenetic distance ratios among all three bacteria and whose corresponding gene was reported (by a ChIP-seq study) to be regulated by RpoN were included in the Venn diagram. (E) Schematic illustration of four classes of RpoN binding sites based on their positions in relation to gene coding sequences. (F) Distribution of each class of RpoN binding site.

We set a cut-off for the PSSM of scores >7 for high-confidence peaks, which yielded 103 peaks encompassing 112 binding sites (Fig. 2C; Table 1 and 2). There were relatively few ChIP-seq peaks without motifs (16 out of 119; Table S2) compared with the findings in some other studies (23, 27, 28). This probably reflects the high-stringency analysis used, which limits the number of false positives commonly found in ChIP data from highly transcribed regions (31, 32). The majority of the peak-associated RpoN binding sites were found in bacteria expressing RpoN-V5 from its native promoter during logarithmic growth. The corresponding samples from bacteria overexpressing RpoN-V5 lacked six of those peaks, and even more peaks were missing from the samples of virulence inducing conditions. The reasons for these differences are not obvious, but in the case of peaks missing in bacteria induced for expression of the virulence plasmid, the availability of exposed sites might have been affected by structural changes in the chromosome *per se* that can be part of mechanisms suppressing chromosomal gene expression (24, 25). In bacteria overexpressing RpoN-V5 there might be a saturation effect at high RpoN-V5 concentrations, with the precipitation of RpoN molecules not associated with any binding site that dilute samples resulting in relatively less precipitated DNA where DNA fragments from low abundant binding sites are missed. Notably, unlike other sigma factors, RpoN can bind its DNA sequence without RNAP, although the binding is 10-fold less efficient than the RpoN:RNAP complex (33). ChIP-seq should therefore also detect RpoN-DNA interactions that are independent of active transcription.

**Table 1:**
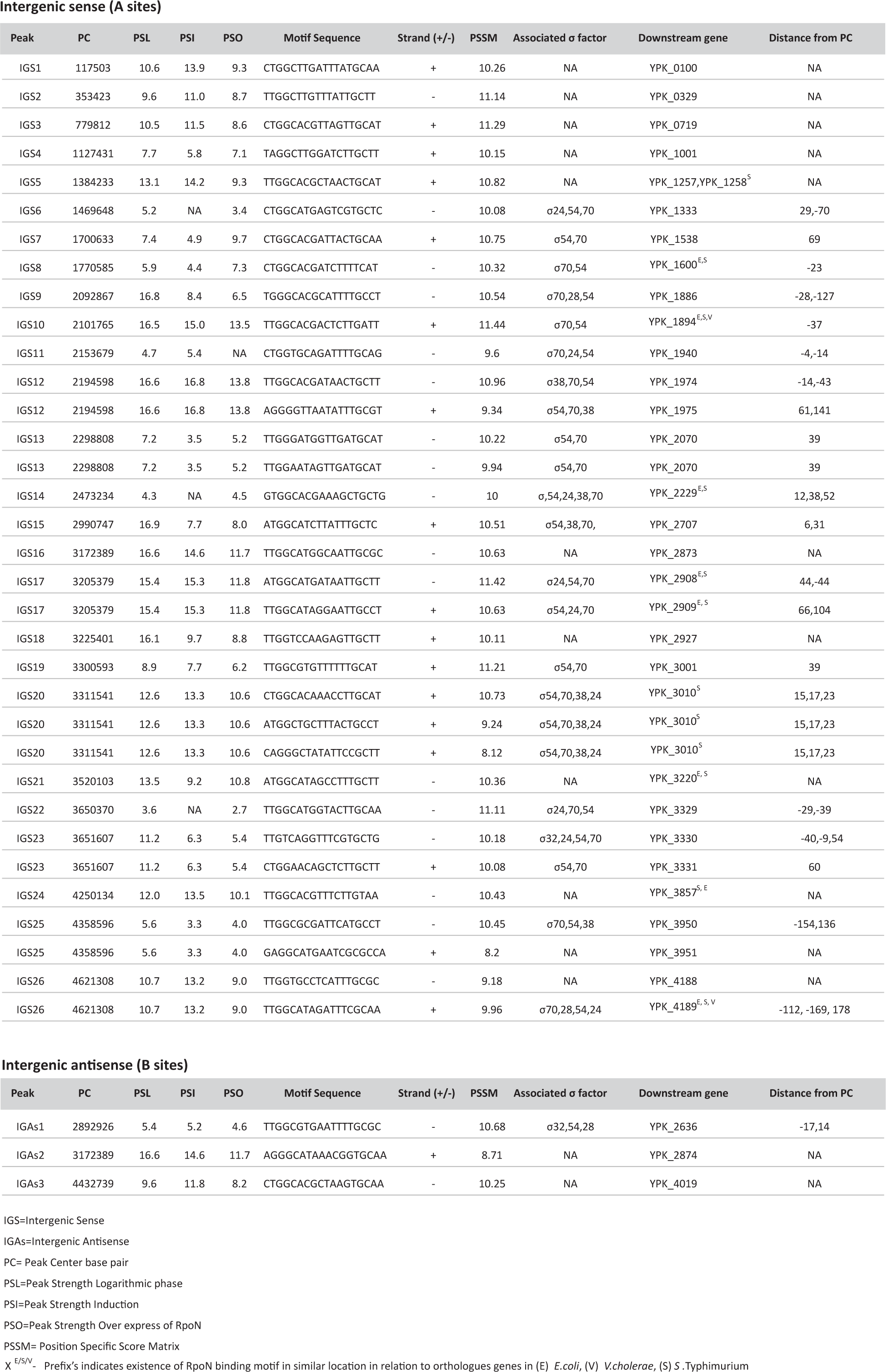
Intergenic sense (A) and antisense (B) RpoN binding sites.

**Table 2:**
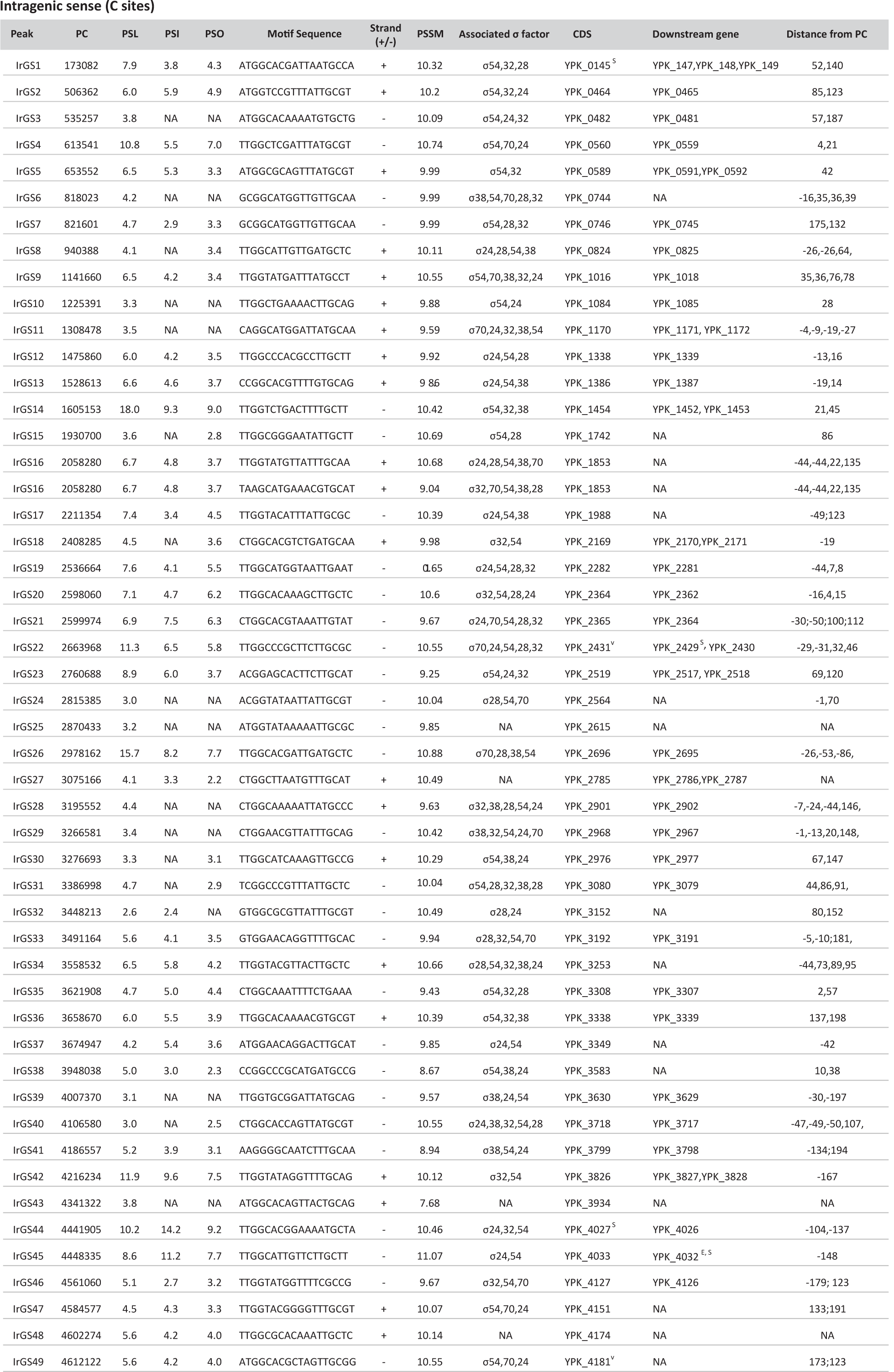
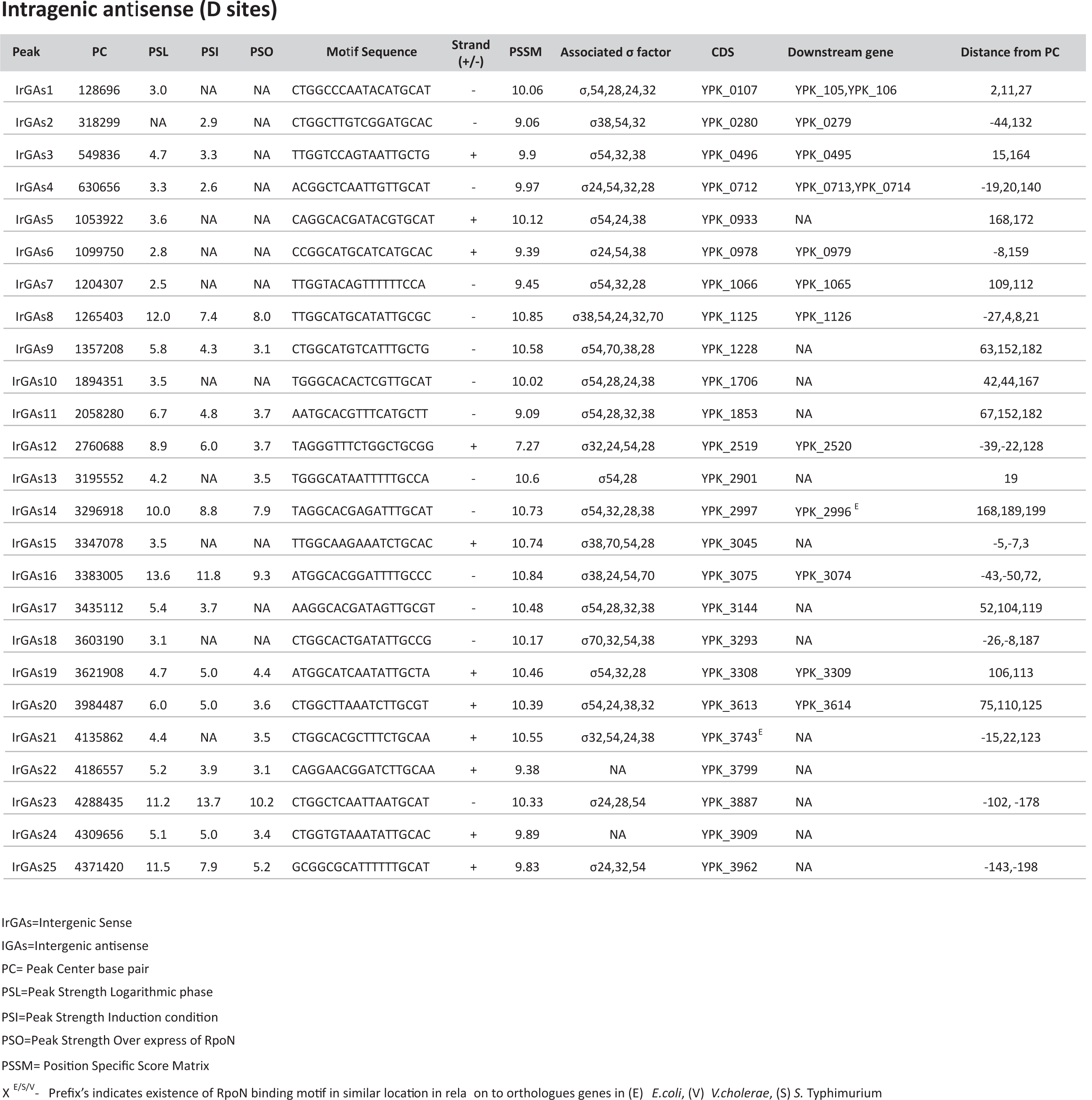
Intragenic sense (C) and antisense (D) RpoN binding sites.

The robustness of the analysis was further verified by the identification of RpoN binding sites at intergenic regions upstream of genes previously shown to be regulated by RpoN in other bacteria. Examples here are YPK_1894 (*pspA*), YPK_3220 (*glnK*), YPK_3857 (*pspG*) and YPK_4189 (*glnA*) (8, 27, 28). Comparing our high-stringency peaks with the peaks identified by ChIP-seq in *E. coli and S*. Typhimurium, it was obvious that the location of many in relation to coding DNA sequences (CDS) was conserved, and some were also shared with *V. cholerae* (Fig. 2D, Table 1, 2, and 3). The conserved locations of RpoN binding sites included locations upstream of *pspA* and *glnA* with orthologs in *E. coli*, and *S*. Typhimurium and *V. cholerae*, for example, YPK_2229, YPK_2908, and YPK_1600, with orthologs in *E. coli*, and *S*. Typhimurium (Table 3). There were also many novel binding sites identified in *Y. pseudotuberculosis*, where a majority were intragenic with some also on the non-coding strand (Fig. 2D, E and F). Presence of sense and antisense intragenic binding sites for RpoN is in accordance with what been previously found for *S*. Typhimurium *and E. coli* (27, 28). The function and mechanisms of RpoN intragenic binding are generally unknown, but there are examples where this binding can drive transcription of downstream genes with long 5’ UTRs (27, 34). The putative RpoN binding sites identified were divided into groups (A–D) based on their position and orientation. Group A (26 sites) and B (3 sites) comprise binding sites in intergenic regions. Group A sites are oriented towards the 5’end of the nearest coding sequence and those in group B are oriented towards the 3’end of the neighboring coding sequence (Fig. 2E and F; Table 1). Groups C (49 sites) and D (25 sites) comprise intragenic binding sites; those in group C are oriented in the sense direction and those in group D in the anti-sense direction (Fig. 2E and F; Table 2). In general, binding sites in group A were associated with stronger peaks than the sites in groups C and D (Table 1 and 2, Fig. S2). Further, compared with the intergenic binding sites among which a relatively large fraction appears to be conserved among *E. coli, S*. Typhimurium, and *Y. pseudotuberculosis*, the fraction of conserved intragenic RpoN-binding sites was considerably lower (Table 1, 2, and 3).

**Table 3:** RpoN binding motif with similar location in relation to orthologues genes.

| Peaks | Locus Tag | Description | Type | *Y. pseudotuberculosis* | *E. coli.* | *V. cholerae* | *S. Typhimurium* |
| --- | --- | --- | --- | --- | --- | --- | --- |
| IGS8 | YPK_1600 | Glutamine ABC transporter periplasmic protein, GlnH | A | ✓ | ✓ |  | ✓ |
| IGS10 | YPK_1894 | Phage shock protein, PspA | A | ✓ | ✓ | ✓ | ✓ |
| IGS14 | YPK_2229 | Bifunctional succinylornithine aminotransferase, AstC | A | ✓ | ✓ |  | ✓ |
| IGS17 | YPK_2908 | Sensor kinase, ZraS | A | ✓ | ✓ |  | ✓ |
| IGS17 | YPK_2909 | Zinc resistance protein, ZraP | A | ✓ | ✓ |  | ✓ |
| IGS21 | YPK_3220 | Nitrogen regulatory protein P-II 2, GlnK | A | ✓ | ✓ |  | ✓ |
| IGS24 | YPK_3857 | Phage shock protein G, PspG | A | ✓ | ✓ |  | ✓ |
| IGS26 | YPK_4189 | Glutamine synthetase, GlnA (Nitrogen metabolism) | A | ✓ | ✓ | ✓ | ✓ |
| IrGS45 | YPK_4032 | Lipopolysaccharide biosynthesis protein, WzzE | C | ✓ | ✓ |  | ✓ |

To reveal possible sigma factor cross talk in *Y. pseudotuberculosis*, we also screened for RpoD, RpoE, RpoS, RpoH, and FliA binding sites close (+/– 200 nt) to the identified RpoN binding sites. This screen showed many potential dual and sometimes triple sigma factor binding regions close to each other in 68% of all A sites. Even more multiple binding regions were found associated with intragenic sites, with clusters of 2–5 sigma factor sites in more than 90% of all C and D sites. Notably, all sigma factor binding regions associated with A sites had a binding site for RpoD (σ70), and nearly half of them contained sites for RpoE (σ24) and/or RpoS (σ38). In contrast, binding sites for FliA (σ28) were very rare and sites for RpoH (σ32) was absent. The multiple binding sites associated with C sites included sites for RpoE (60%), RpoH (54%), RpoS (40%), FliA (42%), and RpoD (30%). D sites were similar, with binding sites for RpoS (64%), RpoH (60%), FliA (52%), RpoE (48%) and RpoD (20%).

### Deletion of *rpoN* has a large impact on the *Y. pseudotuberculosis* transcriptome, with both direct and indirect effects

Next, we aimed to determine whether the identified binding sites could indeed be coupled to gene expression in *Y. pseudotuberculosis*. For this we employed RNA-seq on WT and Δ*rpoN* bacteria at 26°C in stationary phase and at 37°C with virulence induction. Analysis of differentially expressed genes revealed a markedly different gene expression pattern in the Δ*rpoN* strain compared with the isogenic WT strain. More than 500 genes were found to be differentially regulated at 26°C stationary phase: 294 were downregulated and 213 upregulated in Δ*rpoN* (Fig. 3A and 3B). The effect was even more pronounced at 37°C in virulence-inducing conditions: almost 1700 genes were affected, with 766 genes downregulated and 929 upregulated. The reason for this discrepancy, with a much higher number of genes affected during virulence-inducing conditions compared with 26°C stationary phase, might be a higher degree of stress associated with the former. This is a condition known to involve activation of different alternative sigma factors as well as other global regulators. Functional annotation analysis of the differentially expressed genes showed down-regulation of genes involved in nitrogen metabolism, flagellar assembly, chemotaxis, and quorum sensing in both conditions (Fig. 3C and Fig. S3). There was also downregulation of fatty acid biosynthesis and metabolism, but this was not seen in samples of bacteria subjected to virulence-inducing conditions. For these samples, additional pathways were affected by the deletion of *rpoN*, including, for example, low expression of genes involved in T3SS that normally is highly upregulated at this condition, downregulation of DNA replication, and amino acid biosynthesis, and upregulation of genes involved in carbon metabolism, gluconeogenesis, and ribosomal organization, with the latter possibly reflecting the stress of translational reprogramming.

**FIG 3.**
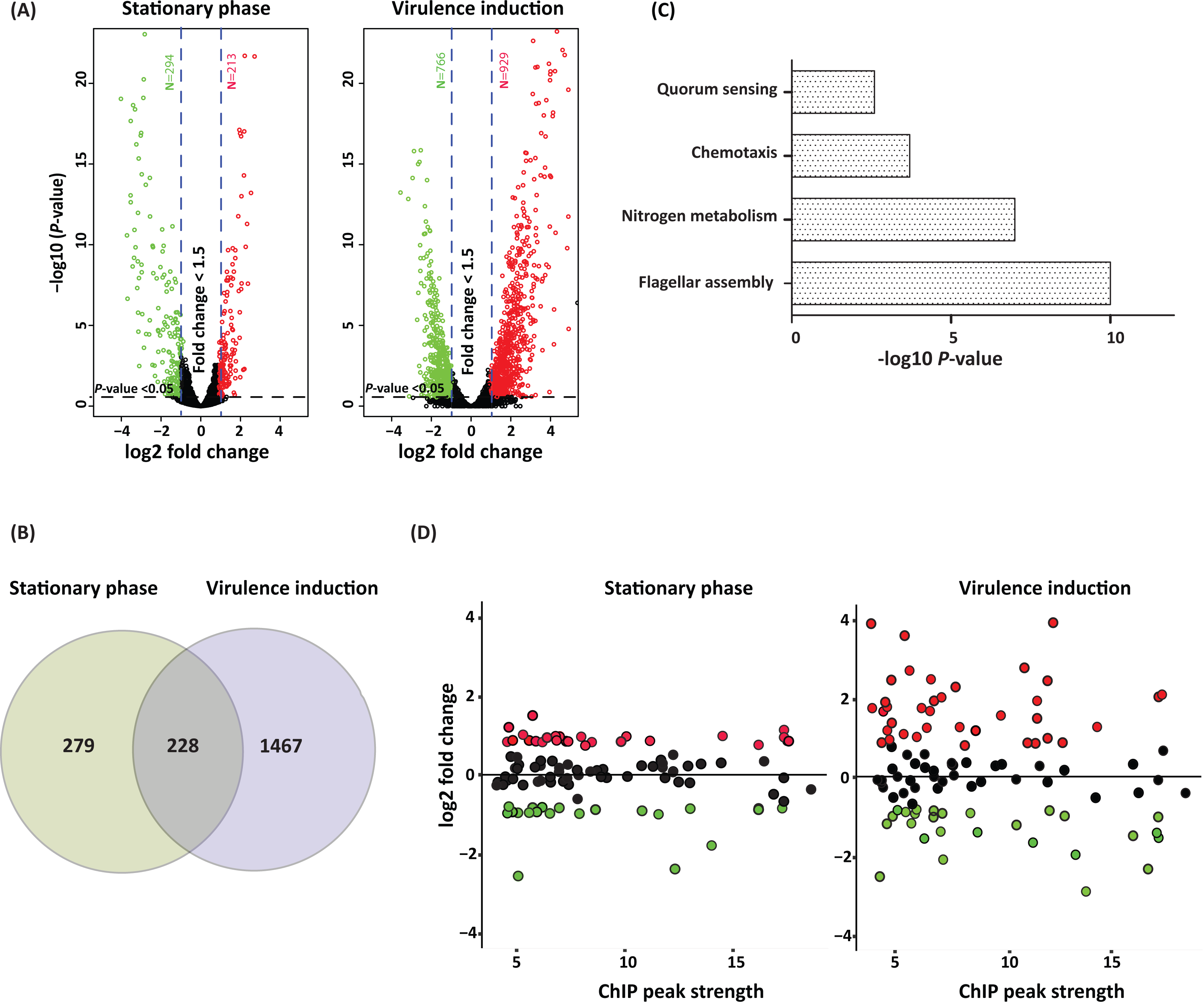
Deletion of *rpoN* has a substantial influence on the *Y. pseudotuberculosis* transcriptome. (A) Volcano plot showing differentially expressed genes in the *Y. pseudotuberculosis* Δ*rpoN* strain compared with the WT strain at stationary phase (OD = 2,0 at 26°C) and during virulence induction (Ca^2+^ depletion, at 37°C). (B) Venn diagram showing the number of differentially expressed genes under the indicated conditions. The criteria for differential expression shown in A and B were fold changes >1.5 with a *P* value <0.05. (C) Pathway enrichment (KEGG) of genes differentially expressed both in stationary phase and after virulence induction. Pathways are ranked by the negative log 10 of the *P* value of the enrichment score. The *P* values were calculated using the Bonferroni correction. (D) Plots showing expression of genes associated with intergenic or intragenic high-confidence peaks. The y-axis indicates expression values from RNA-seq (log2 fold-change of Δ*rpoN*/WT) and the x-axis indicates the strength of the associated ChIP peaks. For both A and D, significantly upregulated (red) and downregulated (green) genes, as well as non-significant changes (black circles) are indicated.

Although the expected effects of RpoN deletion, such as reduced expression of genes involved in nitrogen metabolism, flagella, and quorum sensing were obvious, the effect on the *Y. pseudotuberculosis* transcriptome was huge, clearly reflecting deletion of a global regulator. This accords with previous studies, which found that the absence of RpoN commonly results in a global effect involving both direct and indirect effects on gene expression (9, 23, 35). The differential gene expression analysis revealed changed expression levels of other sigma factors in the Δ*rpoN* strain (Table S3). Expression of RpoD, for example, was significantly upregulated, whereas expression of RpoE was downregulated both in stationary phase and in virulence-inducing conditions. RpoS on the other hand was upregulated in stationary phase but downregulated during inducing conditions. In addition to other sigma factors, the mRNA levels of other transcriptional regulators were affected, such as CpxR, RovA, FlhCD, and others (Table S3). Hence, these effects, together with effects on the transcription of other sigma factors, are expected to contribute extensively to indirect effects on gene expression, adding further complexity to the data set.

### Identified RpoN-binding sites mediates both positive and negative regulation of gene expression

Given the complexity, including indirect effects, in the RNA-seq data set, we next aimed to reveal direct effects of RpoN. The RpoN binding sites identified in the ChIP-seq analysis likely include both active promoters driving transcription and suppressive and silent binding of RpoN to the chromosome. The potential RpoN binding sites identified were therefore matched with detected changes in gene expression levels (Fig. 3D). Among the identified binding sites in group A, 88% were associated with differential gene expression, involving both upregulated and downregulated genes (Fig. S2). The intragenic C and D sites were also associated with differential gene expression, 79% for the C sites and 68% for the D sites. For C sites, the differential gene expression included both upregulation and down-regulation of genes containing the motif as well as downstream genes. For the antisense D sites, there was a larger fraction of upregulated than downregulated genes, suggesting potential suppressive effects of the RpoN binding (Fig. S2). In general, a relatively large portion of the differentially expressed genes associated with RpoN binding showed increased expression in the Δ*rpoN* strain. Binding by RpoN might suppress transcription by nearby sigma factors and possibly other transcription factors binding to the same region, where the absence of RpoN would then allow transcription to occur (8, 27). Also, collision as a consequence of RpoN intragenic binding has been suggested (36, 37).

To explore the potential importance of the RpoN binding sites identified in *Y. pseudotuberculosis*, we set out to mutate some of them to reveal their effects on gene expression. For this we chose binding sites with locations indicative of putative positive or negative regulation by RpoN, which was the case for many A and D sites. We could not identify any C-site ChIP-seq peak indicative of transcriptional activation of downstream genes in our data set. The sites were mutated by the exchange of 3–6 nucleotides in the conserved TGG and TGC sequences of the RpoN binding motif. In intragenic motifs, the exchanged nucleotides were selected in order to minimize changes in the encoded protein (see Table S5 for details). The selected sites included seven A and five D sites (Fig. 4 and Table S4) and the effects of the mutations on gene expression were verified by qPCR. For all putative activating A sites that is, those represented by down-stream genes downregulated in the absence of RpoN point mutations in the RpoN binding motifs in the WT strain resulted in reduced transcription. This class of binding sites also showed the highest degree of conservation. The most prominent effect of the disruptive nucleotide exchange was seen for *pspA* (YPK_1894), a gene known to be activated by RpoN (38, 39). There were also indications of inhibitory effects of RpoN binding to A sites, where the expression of down-stream genes was increased in both Δ*rpoN* and the corresponding binding site mutant. Intriguingly, four of the five binding site mutations in D sites resulted in increased expression of the CDS on the opposite strand, suggesting a suppressive effect of RpoN binding. We also mutated some of the C sites, but here no effect on transcription compared with that in the WT strain could be seen. Notably, peaks associated with C sites were commonly flatter and broader than the peaks covering the mutated A and D binding sites (Fig. S4).

**FIG 4.**
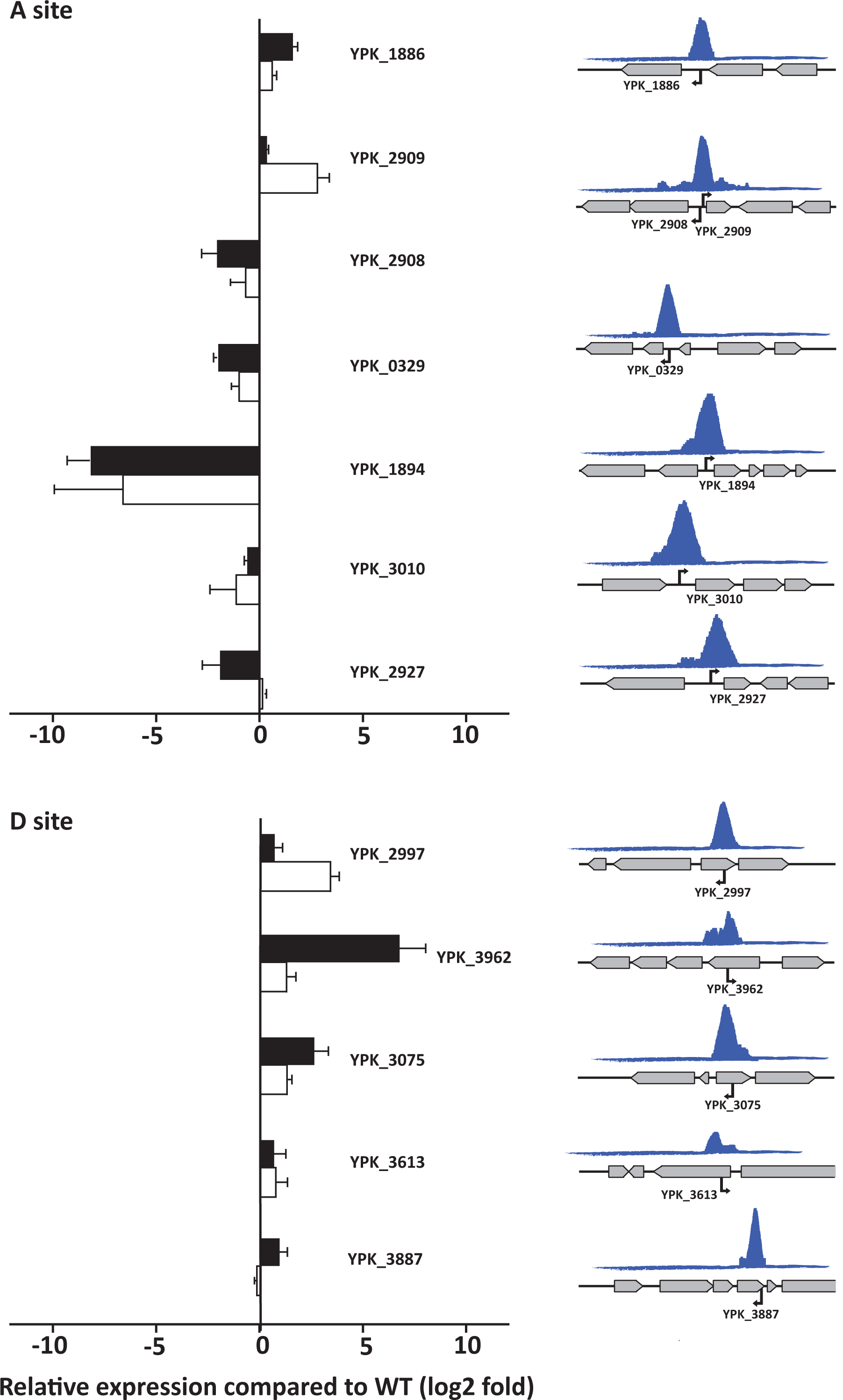
Validation of productive RpoN-binding. (Left) Relative expression of indicated genes in bacteria when the associated RpoN binding site is mutated (white bars) and in the Δ*rpoN* strain (black bars). Expression of genes in the mutant strains was determined by qPCR and presented as log2 fold relative expression compared with expression in the WT strain, where positive values represent suppressed transcription and negative indicate activated transcription. Results are shown for expression during virulence-inducing conditions for YPK_0329, YPK_1886, YPK_1894, YPK_2927, YPK_2997, YPK_3010, YPK_3075, YPK_3613, YPK_3887, and YPK_3962, and for expression during stationary phase at 26°C for YPK_2908 and YPK_2909. Values represent means ± SEM of three independent experiments. (Right) Schematic illustration of the shapes and genomic contexts of ChIP-peaks together with the positions and orientations of RpoN-binding of associated motifs.

Taken together, we have by mutagenesis been able to verify effects of RpoN binding to the binding motifs identified in a ChIP-seq screen of *Y. pseudotuberculosis*. Activating as well as suppressing effects of intergenic RpoN-binding were verified by mutating intergenic A sites. Among these verified productive RpoN-binding sites, some were known, such as those in PspA, PspP, and GlnA, whereas RpoN regulation of TppB, UgpB, and DkgA is described for the first time. There were also indications of inhibitory effects of RpoN binding to intragenic D sites. As discussed earlier, how RpoN suppresses transcription is less clear but for binding to the sense strand it might occur by steric hindrance, either by the RpoN-RNA polymerase complex or RpoN alone binding to the DNA. How this can affect expression from the opposite strand, as would be the case for the observed inhibitory effect of mutating RpoN binding D sites is less clear, but it might involve disturbed strand separation. For intragenic RpoN binding, we saw effects on transcription only by mutating binding sites on the anti-Sense strand. Thus, intragenic binding by RpoN to coding regions on the sense strand is likely silent, and may be used for storage of RpoN-RNAP. RpoN-mediated suppression by binding to internal anti-sense sequences of genes encoded on the opposite strand has not been shown previously and its mechanism remains to be elucidated.

## MATERIALS AND METHODS

### Bacterial strains and growth conditions

Strains and plasmids are listed in Table S5. *Yersinia pseudotuberculosis* strain YPIII was used in this study. *Escherichia coli* S17-1 λpir was used for cloning and conjugation. Antibiotics were used at the following concentrations: ampicillin (100 µg/ml), kanamycin (50 µg/ml), and chloramphenicol (25 µg/ml). Motility was tested on Luria Bertani (LB) medium with 0.6% agar. Biofilm assays were carried out as described previously, using LB medium in glass tubes (40). All strains were routinely grown at 26°C in LB medium containing kanamycin (50 µg/ml). For ChIP- and RNA-Seq analyses, cultures were grown in LB medium to the desired OD_600_. Arabinose (0.005%) was used for 30-min induction at 26°C for overexpression of *rpoN*. To reach the virulence induction condition, overnight bacterial cultures were diluted to OD_600_ 0.05 in LB and grown at 26°C. After 1 h, calcium was depleted by adding 5mM EGTA and 20 mM MgCl_2_ and cultures were shifted to 37°C (26).

### Strain construction

In-frame gene deletion and insertion of the V5 epitope and binding-site mutations in *Y. pseudotuberculosis* were performed using an In-fusionHD cloning kit (Clontech) according to the manufacturer’s instructions. Briefly, the flanking regions of the respective gene were amplified by PCR and cloned into the suicide vector pDM4. This construct was used to transform S17-1, and then transferred into recipient strains through conjugation. Conjugants were purified and incubated on 5% sucrose to recombine out the vector together with wt sequence. Deletion or mutation was confirmed by PCR. For trans-complementation, the gene was PCR amplified and cloned into the pBAD24 plasmid. For gene induction, rpoN and a C-terminal 3×V5 epitope tag were PCR amplified and cloned into the pBAD18 plasmid. All constructs were verified by sequencing. Primers used in this study are listed in Table S6.

### Motility and biofilm assays

Determination of swimming motility was performed as described previously (41). A 5 µl aliquot of a diluted overnight culture (OD_600_ of 1.0) were spotted at the center of a 0.6% LA soft agar plate and incubated at 26°C for 24 h. Bacterial motility was determined by measuring the diameter of the bacterial growth area.

Biofilm formation was determined as previously described (40). Overnight cultures were diluted to an OD_600_ of 0.05 and grown to an OD_600_ of 0.5 in a 26°C shaking water bath. A 1 ml aliquot of the bacterial culture was pelleted and dissolved in 2 ml fresh LB. The suspension was transferred to glass tubes and incubated for 48 hours at 37°C without shaking. After incubation, the bacterial suspension was discarded and tubes were gently washed 3 times with PBS and stained with 0.1% crystal violet (Sigma-Aldrich) for 15 min, followed by 3 successive washes with PBS. The biofilms on the tube surface were thereafter dissolved with 33% acetic acid for 15 min and the absorbance at 590 nm was measured with an Ultrospec 2100 pro spectrophotometer (Amersham Biosciences, Piscataway, NJ).

### RNA isolation and Illumina sequencing

Total RNA was isolated from three independent biological replicates of WT *Y. pseudotuberculosis* and an isogenic rpoN deletion strain at stationary phase and under virulence-inducing conditions. For isolation and purification of RNA, the Trizol method (Ambion Life Technologies, Carlsbad, CA) and the Direct-zol RNA kit (Zymo Research, USA) were used. To remove unwanted DNA, purified RNA was treated with DNAse for 30 min at room temperature according to the manufacturer’s instructions. RNA quantity was measured by Qubit (Nordic Biolabs AB, Sweden). RNA from stationary phase bacteria was collected after 8 hours of growth (OD_600_ ∼2) at 26°C in LB. For virulence-inducing conditions, RNA was isolated from cells grown exponentially at 26°C, shifted to 37°C, and incubated in Ca^2+^-depleted medium for 3 h.

For sequencing, cDNA libraries were prepared using the ScriptSeqTM Complete Kit (Epicentre, Madison, WI, USA) according to the manufacturer’s instructions. Ribosomal RNA was depleted from total RNA using a Ribo-Zero rRNA Removal Kit for Bacteria (Epicenter, Madison, WI, USA) following the manufacturer’s protocol. The resulting cDNA libraries were purified using AMPure XP (Beckman Coulter) and quantified using an Agilent 2100 Bioanalyzer. Sequencing was done using an Illumina MiSeq.

Raw RNA-seq reads were trimmed from 5’ and 3’ ends by Trimmomatic (42) until all the adapter and low-quality bases (Phred quality score >30) were removed and the sequences passed Quality checking by FastQC (43). ProkSeq, a complete RNA-seq data analysis package for prokaryotic was used for further RNA-seq data processing, quality control, and visualization (44). It includes all the tools mentioned below as well as SAMtools (45) and BEDtools (46). Reads were aligned to *Y. pseudotuberculosis* YPIII chromosome (NC_010465) and plasmid (NC_006153) using bowtie2 (47) with the unique-mapping option. Post-mapped read quality checking was done using RseQC (48), and the numbers of reads for each gene were counted using featureCounts (49). Differential gene expression was determined using DEseq2, using the shrinkage estimation of dispersion option (lfcSrink =True) to generate more accurate estimates of differential expression in fold changes (50). Differential expression of a gene was defined using absolute log2 fold change values of ≥1.5 and false-discovery-rate values of <0.05. Figures are plotted using the ggplot2 package in R (linux version 4.0.2) and GraphPad Prism (version 8.0)

### Chromatin immunoprecipitation followed by next generation Sequencing (ChIP-Seq)

*Y. pseudotuberculosis* Δ*rpoN* expressing RpoN::V5 in trans under the inducible araBAD promoter (YPIII, Δ*rpoN*/pIBX, p*rpoN*::V5) as well as *Y. pseudotuberculosis* expressing RpoN:V5 in *cis* under its native promoter (YPIII, *rpoN*::V5/pIBX) were grown overnight in LB, diluted to OD_600_ 0.05, and grown for 4 h at 26°C, at which point the p*rpoN*::V5 plasmid was induced with 0.005% arabinose for 30 min. Virulence-induced cultures were shifted to 37°C after 1 h at 26°C and incubated for 75 min in calcium-depleted conditions.. Non-tagged WT *Y. pseudotuberculosis* grown at 26°C for 4 h was used as a negative control. Cultures were cross-linked with 1% formaldehyde at room temperature for 10 min. For subsequent ChIP-seq, 10 OD_600_ were used for each biological replicate. Chromatin immunoprecipitation was performed as described previously, with slight modifications (27). The 1 ml samples were sonicated in AFA milli-tubes using a Covaris E220 sonicator with the following settings: peak power, 140W; duty factor, –5%; cycles/burst, 200; time, 16 min. Immunoprecipitation was done using 90 μL Anti-V5 Agarose Affinity Gel (Sigma A 7345) overnight at 4°C. ChIP DNA was eluted in 100 μl elution buffer, treated with RNase A and proteinase K, and purified using a ChIP DNA clean & Concentrator Kit (Zymo Research, USA). AMPure XP magnetic beads (Beckman Coulter) 1.5× and 0.8× were used to remove adapters and clean and concentrate the DNA. Failsafe polymerase (Nordic Biolabs AB, Sweden) was used for efficient library amplification.

### ChIP-seq Data analysis

A custom pipeline in Python was used for ChIP-seq data analysis, data visualization, and downstream bioinformatic and computational analyses (the pipeline is available on request). The quality of raw reads generated from Hiseq 2500 was checked using FastQC (43). Raw reads were trimmed by 5 nt at both the 5’ and 3’ ends and aligned with the reference *Y. pseudotuberculosis* YPIII chromosome (NC_010465) and plasmid (NC_006153) using BWA (51). Aligned reads were converted to BAM files using SAMtools (45) and duplicate reads were removed with picard (52) by deduplicate function. Before peak calling, two biological replicates for each sample and the input (control) were merged using SAMtools. Peak calling was done using MACS (2.1.2) (53) with the following custom settings: log2FC>1 over input, --broad –g, --broad-cutoff = 0.1. Identified peak co-ordinates in the genome were used to identify the probable regulated genes using the R packages ChIPpeakAnno and ChIPseeker (54, 55). De novo motif prediction was done using the bioconductor package BCRANK (56). A position-weight matrix (PWM) of the predicted motif was used to map the motif presence around the peak center using the FIMO-meme package (linux version) (57). A region of 50 bases upstream and downstream sequence of the peak center was used as input for searching for the PWM of the motif. The SIGffRid tool was used to predict the binding sites of other sigma factors within 200 bases upstream and downstream of each predicted RpoN-binding peak center. To check the conservation of RpoN-regulated genes, cross-species homology prediction was done by comparing *Y. pseudotuberculosis* YPIII, *V. cholerae* 037, *E. coli* 0157, and *S*. Typhimurium LT2 using OrthologueDB (58). Yersinia genes having orthologs in at least one of the species mentioned above were checked for previous ChIP-seq studies (23, 27, 28). Figures are plotted using the ggplot2 package in R (linux version 4.0.2).

### Ethics statement

Mice were housed and treated in accordance with the Swedish National Board for Laboratory Animals guidelines. All the animal procedures were approved by the Animal Ethics Committee of Umeå University (Dnr A108-10). Mice were allowed for one week to conform to the new environment before the experiments start.

### Mouse infections

Female FVB/N mice (Taconic) 8 weeks old were deprived of food and water for 16 hours before infection. For infection of mice, overnight cultures of the *Y. pseudotuberculosis* strains were suspended in sterilized tap water supplemented with 150 mM NaCl, reaching an approximate colony forming unit (CFU) count of 10^7^ CFU/ml for low-dose infection and 10^8^ CFU/ml for acute infection. Mice were allowed to drink for 6 h. The infection dose was calculated based on viable count and the volume of drinking water supplemented with bacteria that was consumed. Frequent inspections of mice were carried out routinely to ensure no prominent clinical signs were overlooked. Infected mice showing notable clinical signs were euthanized promptly to prevent suffering.

The infections were monitored using IVIS Spectrum (Caliper Life Sciences) routinely every third day until 15 dpi (days post infection), and later every week up to 28 dpi. The mice were anesthetized using the XGI-8 gas anesthesia system (Caliper Life Sciences) and 2.5% IsoFloVet (Orion Pharma, Abbott Laboratories Ltd, Great Britain) in oxygen for initial anesthesia and 0.5% isoflurane in oxygen during IVIS imaging. After infection, some mice were euthanized and dissected to analyze bacterial localization and presence in various organs, including the intestine, mesenteric lymph nodes, liver, and spleen. The organs were imaged using bioluminescence imaging. Living Image software, version 3.1 (Caliper Life Sciences, Inc.) was used for image acquisition and data analysis.

### cDNA preparation and qPCR

To validate the ChIP-Seq and RNA-Seq results, qPCR was performed using qPCRBIO SyGreen Mix (PCR Biosystems) and a BioRad CFX Maestro real-time PCR machine. For RpoN binding-site mutants, bacteria were grown for 2.5 hours at 37°C in inducing conditions in LB (OD_600_ ∼0.6) or at 26°C for 8 hours, and total RNA was extracted as described earlier. Isolated RNA was used as template for cDNA synthesis using a RevertAid First Strand cDNA Synthesis kit (ThermoScientific). Experiments were done in triplicate for each mutant. YPK_0831 was selected as an internal control in order to calculate the relative expression levels of tested genes, using appropriate primers (Table S6).

### Data availability

The RNA-seq and ChIP-seq data files have been deposited in Gene Expression Omnibus (GEO) under accession number GSE155608. All the computer code and pipeline used in this studies are available on request.

## ACKNOWLEDGEMENTS

The work has been supported by funding from Knut and Alice Wallenberg foundation (2016.0063), Swedish research Council (2018-02855), and the Medical faculty at Umea University.

We thank SciLife for sequencing facilities. We would like to acknowledge Allison Churcher at the National Bioinformatics Infrastructure Sweden at SciLifeLab for bioinformatics advice.

Conception or design of the work: F.A.K.M.M, K.N,R.C and M.F; data collection: A.K.M.M, K.N,R.C.,R.N, K.A; data analysis and interpretation: F.A.K.M.M, K.N, A.F. and M.F.; drafting of the article: F.A.K.M.M, K.N and M.F; critical revision and contributions to the article: F.A.K.M.M, K.N, A.F., R.N, K.A. and M.F.; final approval of the version to be published: F.A.K.M.M, K.N, A.F., R.N., R.C, K.A. and M.F.

## FIGURE LEGENDS

**FIG S1 High and low-dose infection of *Y. pseudotuberculosis* WT and ΔrpoN in female FVB/N mice**. (S1A) Mice (one experiment, n = 6) were infected with a mean of 4.2 × 10^8^ CFU/ml. IVIS imaging was employed at each time point to determine if mice were infected. Severely sick WT mice were sacrificed before day 7. The color scale represents pseudocolors of the intensity of bioluminescence, where red is the most intense light emission and blue corresponds to the weakest signal. (S1B) Mice (one experiment :WT, n= 40 and ΔrpoN, n= 20) were infected for 28 days with a mean of 4.2 × 10^6^ CFU/ml. IVIS imaging was used to determine if mice were infected. Mice showing acute clinical signs were sacrified. Symptom-free infection was calculated at 28 days after infection.

**FIG S2 Different classes of ChIP peaks and their associated gene expression (WT/ΔrpoN)**. The y-axis represents the fold-change difference between WT and Δ*rpoN* and the x-axis represents the average peak strength.

**FIG S3 Pathway enrichment in t differentially expressed genes (Δ*rpoN*/WT) at stationary phase and under virulence induction**. The pathway enrichment score was calculated as the probability of a gene’s enrichment in a pathway other than by random chance. Pathways are ranked by the negative log 10 of the *P* value of the enrichment score. The *P* values were calculated using the Bonferroni correction and used to evaluate the significance of the pathway in a particular condition. Up- and downregulation of the pathway was predicted by calculating the Z score of differential expression of the genes in a certain pathway.

**FIG S4 Schematic representation of selected intragenic sense (C) RpoN binding sites for qPCR**. The arrow indicates the position and direction of the RpoN binding motif. ChIP peaks are shown in blue, representing their coverage in the genome in logarithmic phase (26°C).

